# Exercise improved P2Y12-regulated microglial dynamics during stroke via endocannabinoid signaling

**DOI:** 10.1101/2023.04.24.538065

**Authors:** Xiao-fei He, Yun Zhao, Ge Li, Jing Luo, Jing-hui Xu, Hai-qing Zheng, Li-ying Zhang, Xi-quan Hu

## Abstract

Microglia are vigilant housekeepers in the adult brain, they continually extend and retract their processes to survey their microenvironment. In a dependent manner involving P2Y12 receptors, microglia undergo morphological and functional changes to form microglia-neuron contacts to protect neurons from damage. By using *in vivo* two-photon imaging, we found that physical exercise (PE) upregulated microglial P2Y12 expression, increased microglial dynamics, and promoted the microglia contacting with neurons in a mouse model of transit middle cerebral artery occlusion (tMCAO). As a result, microglial processes inhibited neuronal calcium overloads, protected against damage of the neuronal mitochondria and synaptic structure. Inhibition of P2Y12 by PSB0739 abolished the protection induced by PE. Mechanistically, we found PE increased the cannabinoid receptor 2 (CBR2) in microglia, and administration of AM630, a CBR2 antagonist, decreased P2Y12R expression and abolished PE-mediated effects. These findings identified endocannabinoid signaling may as the critical regulator of the PE-induced P2Y12-mediated effect whereby PE increased the endocannabinoid system to regulate purinergic signaling, further inducing microglial processes at microglia-neuron contacts to monitor and protect neuronal functioning.

## Introduction

Occlusion of the middle cerebral artery (MCAo) is the most frequent type of ischemia stroke, which may produce paralysis on one side of the body and lead to long-term morbidity and mortality in developed and developing countries^[1,2]^. The ischemic core experiences a sudden reduction of blood flow, with cell death occurring irreversibly in the first hours, but apoptosis in the ischemic penumbra may occur after several hours or days^[2]^. Elevation of the intracellular calcium (Ca^2+^) concentrations critically induces glutamate release from excitatory neurons^[3]^, which is one of the key molecular events initiating apoptosis^[2,4–6]^. However, neuroprotection of antagonists of the pivotal targets of excitotoxicity for ischemic stroke has shown poor results in clinical trials, although Ca^2+^ overload agents that block intracellular Ca^2+^ overload have been investigated as protective agents during cerebral ischemia^[7]^. It is therefore important to identify new therapeutic candidates to inhibit the overload of neuronal Ca^2+^.

P2Y12 is a chemoreceptor for adenosine diphosphate (ADP) that belongs to the Gi class of a group of G protein-coupled (GPCR) purinergic receptors. Microglia continuously extend and retract their processes to produce P2Y12-mediated microglia-neuron somatic contacts^[8–10]^. These contacts have been reported to inhibit neuronal Ca^2+^ overload and protect against neuronal damage from ischemic stroke^[10,11]^. Microglial processes contacting neurons have been shown to be protective as early as 4 h after brain injury^[10]^, but salvageable neurons in the penumbra may be metabolically active up to 17 h following a stroke^[10,12,13]^. It is therefore beneficial to fully understand the protective mechanism of P2Y12-mediated microglial processes during ischemic stroke, to help identify new therapeutic candidates to strengthen the microglial process.

Physical exercise (PE) is a lifestyle choice that increases physical and mental health throughout life^[14]^; it downregulates the expressions of pro-inflammatory cytokines and shifts the microglia phenotype towards neuroprotection^[15]^, to further alleviate the pathogenesis of stroke^[16]^. Nevertheless, the mechanism by which exercise regulates microglial activities and affects microglial extension/outgrowth is not fully known. To this end, we investigated P2Y12 as a putative link between exercise and microglial protection after ischemic stroke. We found that exercise-upregulated microglial P2Y12 expression increased microglia-neuron junctions, improved microglial dynamics, inhibited neuronal Ca^2+^ overloads, and protected mitochondrial homeostasis in intact brains, using *in vivo* two-photon microscopy. However, blockade of P2Y12 by PSB0739 neutralized these protective effects induced by exercise, confirming that exercise enhanced microglial protection in a P2Y12-dependent manner.

P2Y12 has been reported to be specifically expressed on microglia in the brain ^[17]^, although how exercise controls P2Y12 expression as well as P2Y12-dependent microglial outgrowth/retraction, is not fully known. Recent studies reported that endocannabinoid signaling (ECS) increased by physical exercise retained the phagocytic potential of microglia and indirectly altered the neuron-microglia communication system^[18–21]^, making them likely candidates to affect the P2Y12-related microglial processes in future therapeutics. In this study, we identified a function of the endocannabinoid system in microglial activity. We also showed that the endocannabinoid system was highly upregulated during exercise, which protected neurons from excitement-induced toxic brain injury, by inducing microglia-neuron somatic junctions via the expression of CBR2-upregulated P2Y12.

## Methods

### Animals

This study was approved by the Animal Research Committee of the Animal Monitoring Institute of Guangdong Province (Guangzhou, China). All mice used in this study were male at 6–8 weeks of age. CX3CR1-GFP mice were obtained from the Jackson Laboratory (Bar Harbor, ME, USA; B6.129P2(Cg)-*Cx3cr1tm1Litt*/J, catalog number: 005582) and bred in the Laboratory Animal Monitoring Institute of Guangdong Province. C57BL/6J mice were provided by the Laboratory Animal Monitoring Institute of Guangdong Province. The mice were housed under a 12:12 h light: dark cycle (light on from 07:00−19:00 h), with controlled temperature and humidity and food and water provided ad libitum.

### Experimental stroke

Transient middle cerebral artery occlusion (tMCAO) was performed as described by Szalay^[22]^. In brief, mice were anesthetized with isoflurane delivered in a mixture of 30% O_2_ and 70% N_2_O, which was controlled by an anesthesia apparatus. For this purpose, an incision was made in the midline neck region and the common carotid artery, using a silicone-coated monofilament (210−230 µm tip diameter; Doccol, Sharon, MA, USA) inserted into the right internal carotid artery. After 50 min of occlusion, the filament was removed. For the survival period, the mice were kept in their home cages with facilitated access to water and food. The body temperature was maintained throughout surgery using a heating pad. Mice that died during the surgery were excluded from the study.

### Voluntary wheel training and pharmacological treatments

Animals in the exercise group were housed in polypropylene cages with a 16 cm diameter running wheel, which rotated when a mouse climbed onto the wheel^[23]^. The mice in the stationary group were housed in polypropylene cages of the same size (36 cm L × 20 cm W × 14 cm H). The mice were housed under these conditions for 6 weeks before induction of the ischemia-reperfusion model^[23,24]^.

To study the effects of P2Y12 receptor (P2Y12R) antagonists on experimental stroke, animals in the PE+PSB0739 group were subjected to intracerebroventricular injection (i.c.v) with a single dose of PSB0739 ((Tocris, R&D Systems, Minneapolis, USA, 15 µg dissolved in saline) for 24 hour prior to tMCAO ^[10]^. To study the involvement of endocannabinoid signaling in PE-related protection, mice in the URB597 group were subjected to intraperitoneal injection with 1 mg/kg URB597 ^[25]^ (Cayman Chemicals, Ann Abor, MI, USA), and mice in the PE+AM630 group were subjected to intraperitoneal injection with 1 mg/kg AM630 ^[26]^ at 6 weeks before induction of the ischemia-reperfusion model (Fig. 5A).

**Figure 5.**
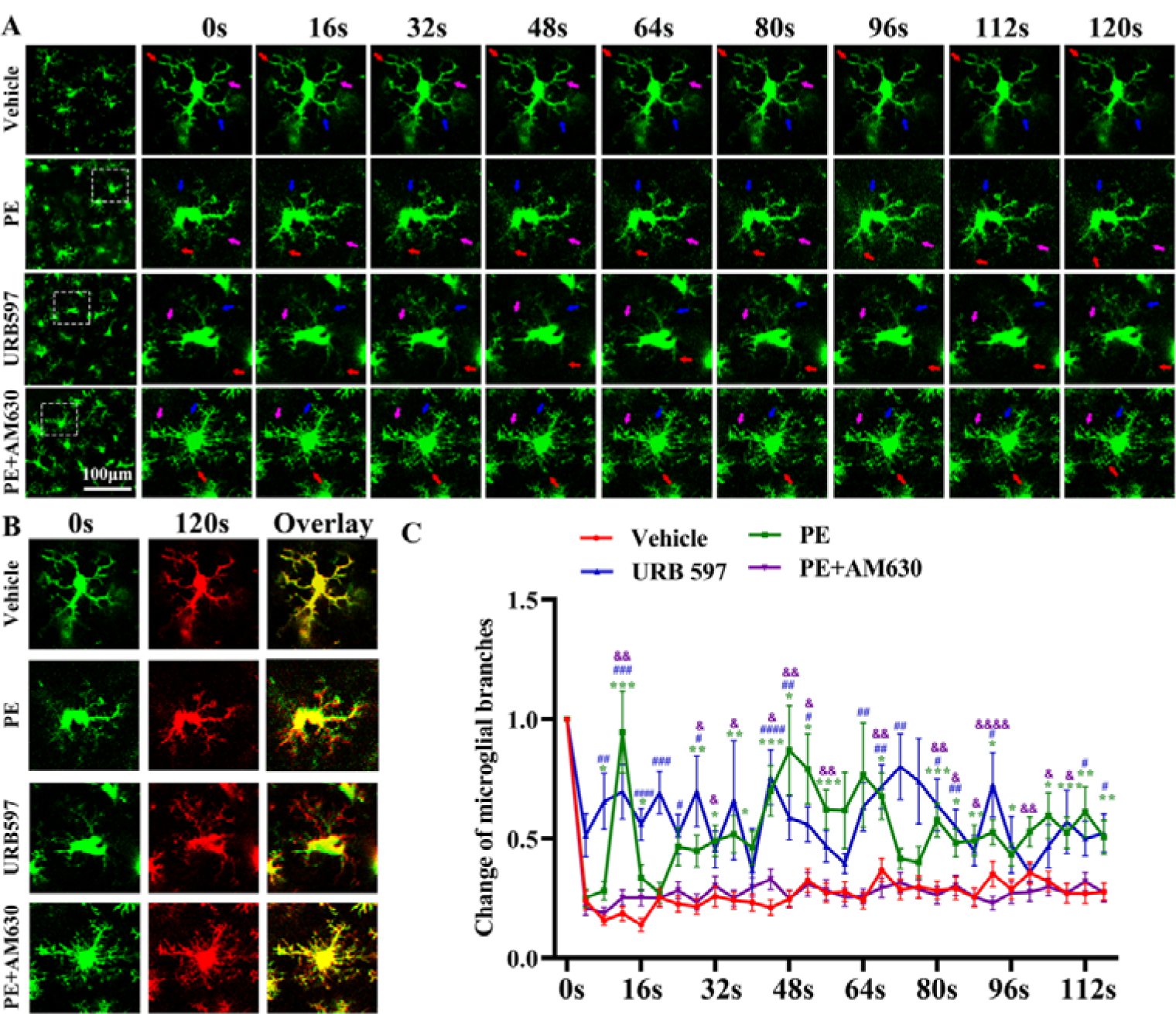
URB597 treatment mimicked the microglial dynamics enhanced by physical exercise (PE), while AM630 decreased the effect of PE. A. Representative graphs showing microglial process extensions or withdrawals in Cx3Cr1-GFP mice (Colorful arrows indicate the microglial branches). B. Sample color-coded image of microglial chemotaxis at the beginning (green) and 2 min (red) after the start of the experiment. C. Comparisons about the change ratios of microglial branches at different time point (*Vehicle *vs.* PE groups; #URB597 *vs.* Vehicle groups; $ PE+AM630 vs. PE groups; Each dataset is expressed as the mean ± S.EM. n=3 branches/microglia, n=3 microglia/mice, n=3 mice/group, ^*,#,$^*P* ≤ 0.05; ^**,##,$$^*P* ≤ 0.01; ^***,###,$$$^*P* ≤ 0.001; ^****,####,$$$$^*P* ≤ 0.0001. n = 4 mice).

### In vivo two-photon imaging of microglial dynamics and neuronal calcium activity

Due to the high transduction efficiency and low immunogenicity of adeno-associated virus (AAV) vectors^[27]^, we intracerebrally injected the AAV-hSyn1-GCaMP6s-P2A-nls-dTomato (Jikai’en Investment, Shanghai, China) in the cortex, 6 weeks before imaging to determine the Ca^2+^ transients and neuronal activity in the peri-infarct area as previously described^[28]^. Because of a high signal-to-noise and ultrasensitive fluorescent indicators, GCaMP6s efficiently detected the increases in intracellular Ca^2+^ under the control of the neuron-specific hSyn1 promoter^[27,29,30]^. Two-photon imaging in the anesthetized mouse was performed using a 2 mm diameter thin cranial window, videos were collected using a Leica SP5 Multiphoton microscope with a coherent laser (512 × 512 pixel) and analyzed using the Leica Application Suite X software (Leica, Wetzlar, Germany)^[24]^. The fluorophore labeling of dTomato in cells expressing GCaMP6s was visualized by excitation at 920 nm and the emitted fluorescence was collected in the red channel, which was defined as regions of interest (ROI). Intracellular Ca^2+^ transients in individual neurons were calculated as the percentage changes from baseline (ΔF/F0) by subtracting this baseline from each point and then dividing by the baseline value. Calcium transient frequency was expressed as the number of fluorescence changes in 1 min (60 s). Twenty cells located 200−300 μm below the pial surface surrounding the clotting site were monitored in each animal. To assess the microglial dynamics, overall changes in lengths for representative processes within single microglia during a 2-min time period were measured, and changes ratio were calculated as length from different time point (t_n_) by subtracting the length at starting point (t_0_) from each point and then dividing by the baseline value [(Legth_t(n)_-Legth_t(0)_)/Legth_t(0)_].

### Histology

Mice were perfused with 50 mL ice-cold saline and 50 mL 4% (w/v) paraformaldehyde in phosphate-buffered saline (PBS; pH 7.4). Coronal brain slices (10μm or 40 μm thick) were sectioned using a frozen microtome (Leica). For Nissl staining, sections were stained with cresyl violet and inspected microscopically. Infarct volume was calculated as described by Denes et al.^[31]^. For immunofluorescence staining, sections were boiled in citric acid buffer (pH 6.0) for 5 min in a microwave oven, then were treated with 0.3% Triton X-100 and 10% goat serum for 1 h at room temperature. The sections were then incubated overnight with primary antibody [1:100 Mouse Anti-Neun, MAB377, Millipore, Massachusetts, USA; 1:100 Chicken Anti-Neun, ab134014, Cambridge, England; 1:400 Rbbit anti-ionized calcium binding adapter molecule 1 (Iba1), Wako, Tokyo, Japan; 1:100 Mouse anti-P2Y12, ab233760, Cambridge, England; 1:100 Rabbit anti-TOMM20, ab186735, Cambridge, England; 1:100 Rabbit anti-Cannabinoid Receptor I, ab259323, Cambridge, England; 1:100 Mouse anti-cannabinoid receptor 2, H00001269-M01, Novus, Colorado, USA; 1:100 Rabit Anti-Synaptophysin, ab32127, Cambridge, England], and then with a secondary antibody at room temperature for 1 h. Apoptosis was detected using a TUNEL Kit (TUNEL Apoptosis Detection Kit; KeyGEN BioTECH, Nanjing, China) according to the manufacturer’s protocol. Images were obtained with × 63 magnification (1024 × 1024 pixels) using a Confocal Microscope (Nikon Eclipse Ni-E, Nikon, Japan).

### Western blotting

Mice were perfused with ice-cold PBS, and tissues in the peri-infarct area were homogenized in 500 μL 1× lysis buffer using a Precellys homogenizer. Total protein levels were quantified using a Pierce™ Microplate BCA Protein Assay Kit (Thermo Fisher Scientific, Waltham, MA, USA). A total of 30 μg protein was separated by SDS-PAGE at 200 V for 45 min using a 4%–12% precast polyacrylamide gel (Novex; Invitrogen, Carlsbad, CA, USA), then transferred to polyvinylidene fluoride membranes (Millipore) at 300 mA for 1.5 h. Membranes were blocked in 5% nonfat skim milk (R&D Systems) for 1 h at room temperature, then the membranes were incubated with the primary antibodies (Rabbit anti-P2Y12, DF10263, Affinity Biosciences, Beijing, China; Rabbit Anti-TOMM20, ab186735, Abcam, Cambridge, England; Rabbit anti-Cannabinoid Receptor I, ab259323, Abcam, Cambridge, England; Mouse anti-cannabinoid receptor 2, H00001269-M01, Novus, Colorado, USA; Rabbit anti-Tubulin beta, AF7011, Affinity Biosciences, Beijing, China; Rabbit anti-iNOS, AF0199, Affinity Biosciences, Beijing, China; Rabbit anti-TNF alpha, AF7014, Affinity Biosciences, Beijing, China; Rabbit anti-IL 10, DF6894, Affinity Biosciences, Beijing, China; Rabbit anti-Beta Actin, AF7018, Affinity Biosciences, Beijing, China; Rabbit anti-GAPDH, AF7021, Affinity Biosciences, Beijing, China) overnight at 4°C. Membranes were then incubated with secondary antibodies. Target protein bands were visualized using a chemiluminescence imaging system.

### Statistics analysis and reproducibility

ImageJ (National Institutes of Health, Bethesda, MD, USA) was used to analyze the data. Three fields in every three sections was selected with a random start, and a total number of nine fields for each mouse were used for counting of cell numbers, and section were analyzed by an experimenter blinded to the experimental condition. SPSS statistical software for Windows, version 19.0 (SPSS, Chicago, IL, USA) or Prism 8.0 software (GraphPad, La Jolla, CA, USA) were used for statistical calculations. Two-way analysis of variance with a Tukey’s post-hoc test for multiple comparisons was used to analyze the microglial dynamics, and one-way analysis of variance with a Tukey’s post-hoc test for multiple comparisons was used to analyze other data. All data are expressed as the mean ± standard deviation of the means (SD or SEM) and a *P* value *<* 0.05 was considered statistically significant. Data are representative of six independent experiments.

## Results

### PE improved microglial dynamics and inhibited neuronal calcium overload in peri-infarct areas in a P2Y12-dependent manner

To explore the mechanism how physical exercise regulates microglial activities and affects microglial extension/outgrowth, we firstly measured the microglial motility and neuronal calcium activity *in vivo* using CX3CR1-GFP transgenic mice to study the effects of PE on microglial processes as well as on neuronal activity at 24 h after the induction of stroke (Fig. 1A). Ultrasound Doppler was performed to validate the ischemic model (Fig. 1B), the imaging site was determined by the two-photon imaging, at the center of an area with a 50% reduction of blood flow (Fig. 1C). Microglia process motility was quantitated in the same cells, showing that the motilities of microglial branches were more pronounced in the PE group (Supplementary vedio.2) than in the vehicle group (Supplementary vedio. 1), which was blocked by treatment with PSB0739 (Supplementary vedio. 3), the individual processes displayed a constant length over time in the vehicle and PE+PSB0739 groups (Fig. 1D-F). Specifically, we measured overall changes in lengths for three representative processes within single microglia during a 2 min time period (Fig. 1D). In vehicle and PE+PSB0739 groups, the microglial images captured at 120s of scanning were overlaid with the same field that was captured at the start of scanning, whereas it showed dynamic state in the PE group (Fig. 1E). Two-way ANOVA results shows that the change ratios of the branch length in PE group were significantly augmented than vehicle group at different time points, which were abolished in PE+PSB0739 group (Fig. 1F).

**Figure 1.**
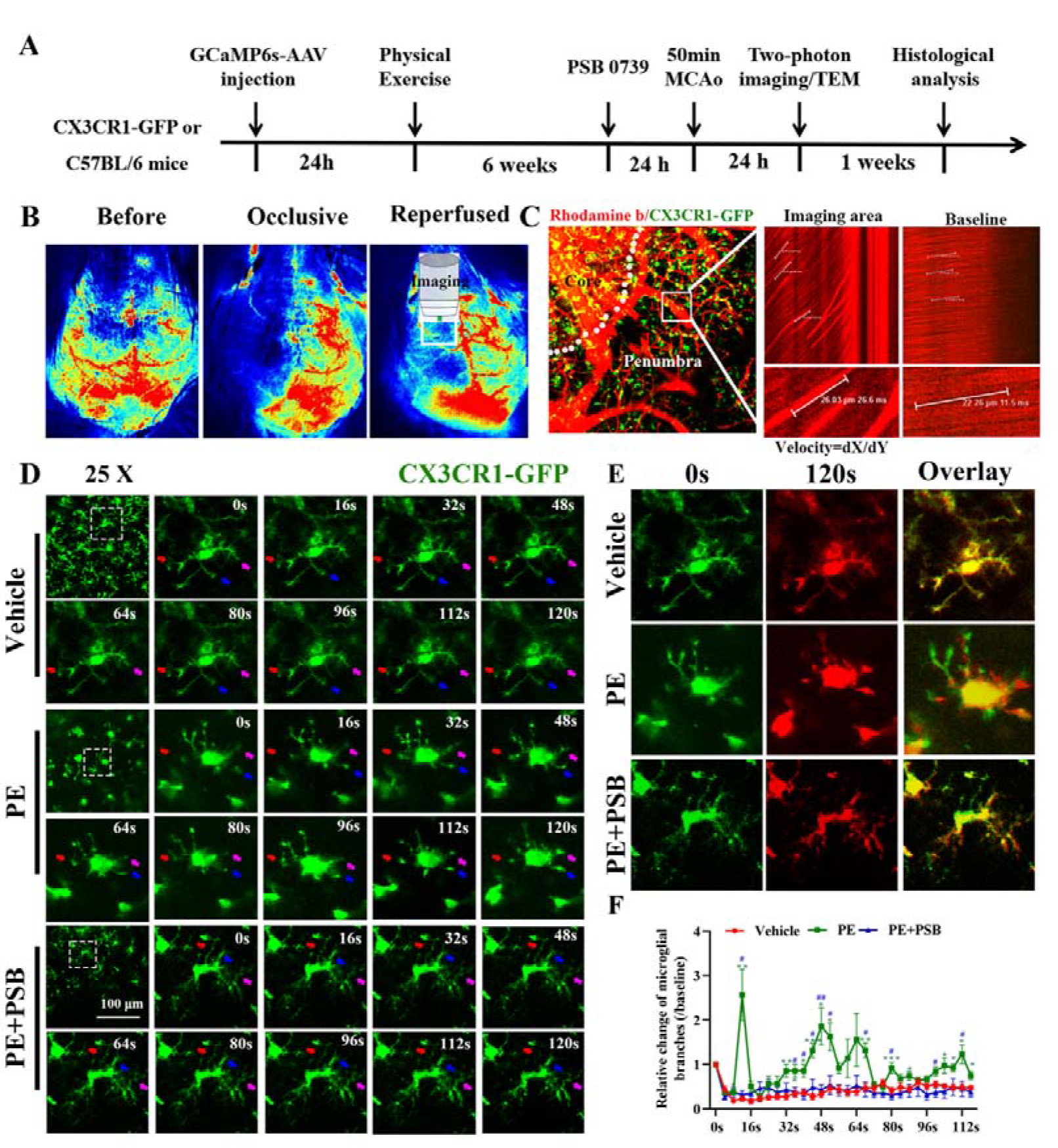
Physical exercise (PE) increased the microglial dynamics in a P2Y12-dependent manner. A. Schematic diagram of time points about this study. B. The representative images of ultrasonic doppler experiment, validating the success of ischemic stroke model and indicating the site of two-photon imaging. C. Representative images using two-photon microscope showing the site and the schematic diagram for blood flow measurement. D. Representative images of GFP-positive microglia at different time points under two-photon microscope, showing the microglial extension or withdrawal in PE group was enhanced compared with that in vehicle group, whereas which was abolished by PSB0739 treatment. E. Comparisons about the change ratios of microglial branch length (Colorful arrows indicate the branches) (3 branches/microglia, 3 microglia/mice, 3 mice/group, Each dataset is expressed as the mean ± S.E.M, \**P* ≤ 0.05; \*\**P* ≤ 0.01; \*\*\**P* ≤ 0.001; \*\*\*\**P* ≤ 0.0001. *PE *vs.* Vehicle groups; **#**PE+PSB0739 *vs.* PE group).

Microglial processes have been reported to monitor neuronal calcium (Ca^2+^) and protect against mitochondrial damage. To characterize the involvement of PE-regulated microglial processes in neuronal Ca^2+^ activity and mitochondrial functioning after stroke, AAV-hSyn1-GCaMP6s-P2A-nls-dTomato was injected into the cortex to detect the activity of neuronal Ca^2+^. Two-photon Ca^2+^imaging showed that the amplitudes of spontaneous intracellular Ca^2+^ transients in the PE group was significantly decreased, when compared with that of the vehicle group (*P* < 0.01), but was not significantly decreased, when compared with that of the PE+PSB0739 group (*P* > 0.05) (Fig. 2A,B). PSB0739 treatment increased the amplitudes of intracellular Ca^2+^ transients, when compared with that of the PE group (*P* < 0.05). In addition, compared with that of the vehicle group, the frequencies of these Ca^2+^ transients were also significantly decreased in the PE group (*P* < 0.001), but not in the PE+PSB0739 group (*P* > 0.05), and the frequency in the PE+PSB0739 group was significantly increased, when compared with that of the PE group (*P* < 0.05) (Fig. 2A,B). Collectively, these findings indicated that microglia made contact with neurons by extending and retracting their branches, which protected against neuronal Ca^2+^ overloads in the peri-infarct areas, and could be increased by PE. However, PSB0739 treatment blocked PE-related microglial dynamics and increased neuronal Ca^2+^ hyperactivity.

**Figure 2.**
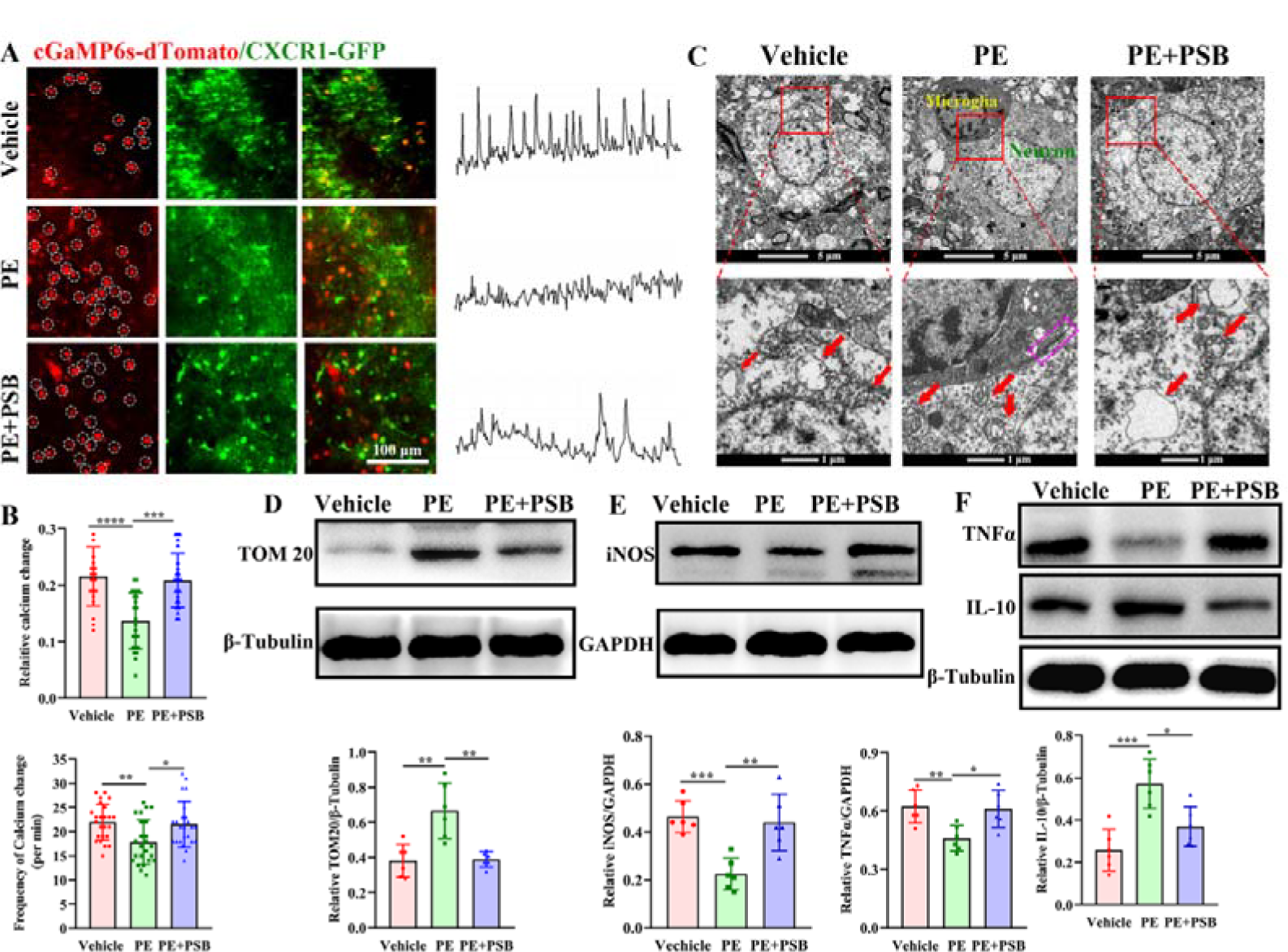
Microglia interacted with neurons, inhibited neuronal calcium hyperactivity, and promoted neuronal mitochondria. A. Left panel: Representative images showing GCaMP6s signal changes in neurons imaged at 24 h after transient middle cerebral artery occlusion. Right panel: Representative images about the relative immunofluorescent intensity changes of GCaMP6s-dTomato. B. Comparisons of the amplitudes and frequencies of GCaMP6s signal changes in the vehicle, PE, and PE+PSB0739 groups. C. Representative images of the neuronal mitochondria and the microglia-neuron membrane junction captured using transmission electron microscope (TEM). Upper panel: images captured using 1200X objective, red box labeled the magnified site showed in the below panel. below panel: images captured using 11000X objective, red arrows indicates the neuronal mitochondria, purple box in PE group indicates the microglia-neuron junction. D. Comparisons of the relative expression of TOM20 among the vehicle, PE, and PE+PSB0739 groups. E. Comparisons of the relative expression of iNOS among the vehicle, PE, and PE+PSB0739 groups. F. Comparisons of the relative expression of TNF α and IL-10 among the vehicle, PE, and PE+PSB0739 groups. Each dataset is expressed as the mean ± SD. \**P* ≤ 0.05; \*\**P* ≤ 0.01; \*\*\**P* ≤ 0.001; \*\*\*\**P* ≤ 0.0001. n = 6 mice.

Mitochondrial-associated membranes are considered to be key integrators of metabolic and immunological imbalances, which determines cellular fates [32]. To investigated the ultrastructural feature of microglia-neuron contacts as well as their protection on neuronal mitochondria, we performed transmission electron microscopy (TEM) (Fig. 2C). In the vehicle groups, microglia-neuron contacts were lacking and the neuronal mitochondria swelled and cristae cracked. However, in PE group, microglia-neuron contacts were composed of reticular membrane structures, intracellular tethers and associated vesicle-like membrane structure within the neuronal body, and the mitochondrial tubular networks and integrity were undamaged. However, this morphological features were not observed in PE+PSB0739 group, and administration of PSB0739 abolished the improvement of PE on the ultrastructue of mitochondria. In a similar manner, in comparisons with the vehicle group, western blots showed that the level of TOMM20 was significantly increased in the PE group, when compared with that of the vehicle group (*P* < 0.001). However, in the PE+PSB0739 group (*P* > 0.05), the expression of TOMM20 in the PE+PSB0739 group was much less than that in the PE group (*P* < 0.05) (Fig. 2D). PE also significantly decreased the levels of pro-inflammatory cytokines including iNOS and TNFα, when compared with that of the vehicle group (Both *P* < 0.01), but there was no difference between the PE+PSB0739 and vehicle groups (Both *P* > 0.05) (Fig. 2E, F). Furthermore, the levels of iNOS and TNFα were both increased in the PE+PSB0739 group, when compared with that of the PE group (*P* < 0.05, *P* < 0.01, respectively) (Fig. 2E, F). In contrast, PE significantly increased the expression of the anti-inflammatory cytokine, IL-10, when compared with that of the vehicle group (*P* < 0.01), which showed no difference between the PE+PSB0739 and vehicle groups (*P* < 0.05), while the level of IL-10 was significantly decreased in the PE+PSB0739 group, when compared with that of the PE group (*P* < 0.01) (Fig. 2F).

### PE protected against the neuronal damge after ischemic stroke

As assessed by immunofluorescence staining, the number of neurons in the peri-infarct area was significantly increased in the PE group (*P* < 0.05), but not in PE+PSB0739 groups (*P* > 0.05) compared to the vehicle group, which showed significantly decreased in PE+PSB039 group, when compared with the PE group (*P* < 0.01). Conversely, the numbers of microglia were significantly decreased in the PE (*P* < 0.01) but not in PE+PSB0739 groups (*P* > 0.05), when compared with the vehicle group, the number of microglia in the PE+PSB0739 group was greater than that in the PE group (*P* < 0.01) (Fig. 3A, B). Western blotting showed that PE significantly increased P2Y12 levels, when compared with that of the vehicle group (*P* < 0.01, PE *vs.* the vehicle groups), and the upregulated P2Y12 was significantly abolished by PSB0739 (*P* < 0.01, PE+PSB0739 *vs.* PE groups), which showed no significant difference between the PE+PSB0739 and vehicle groups (*P* > 0.05) (Fig. 3C). As confirmed by the co-localization immunofluorescence staining, P2Y12 proteins eleavated by PE were mainly located in microglia (Fig. 3D). Overall, these results indicated that PE increased P2Y12 expression, decreased microglial activation and protected against neuronal survival. However, PSB0739 decreased P2Y12 expression and further blocked the PE-associated protection. Additionally, PE significantly decreased the number of TUNEL positive cells in peri-infarct areas, when compared with that of the vehicle group (*P* < 0.01), whereas it showed no difference between the PE+PSB0739 and vehicle groups (*P* > 0.05) (Fig. 3E). Furthermore, compared to the vehicle and PE+PSB0739 groups, the TEM results showed the number of synaptic vesicles were much more and the synaptic membrane structures were clearly visible in PE group (Fig. 3F). Western blots confirmed that PE upregulated the synaptophysin (*P* < 0.05, PE *vs.* the vehicle groups), which was abolished by PSB0739 (*P* < 0.05, PE+PSB0739 *vs.* PE groups). Together, these results indicated that PE protected against mitochondrial and synaptic damage from ischemic stroke, which was blocked by treatment with PSB0739.

**Figure 3.**
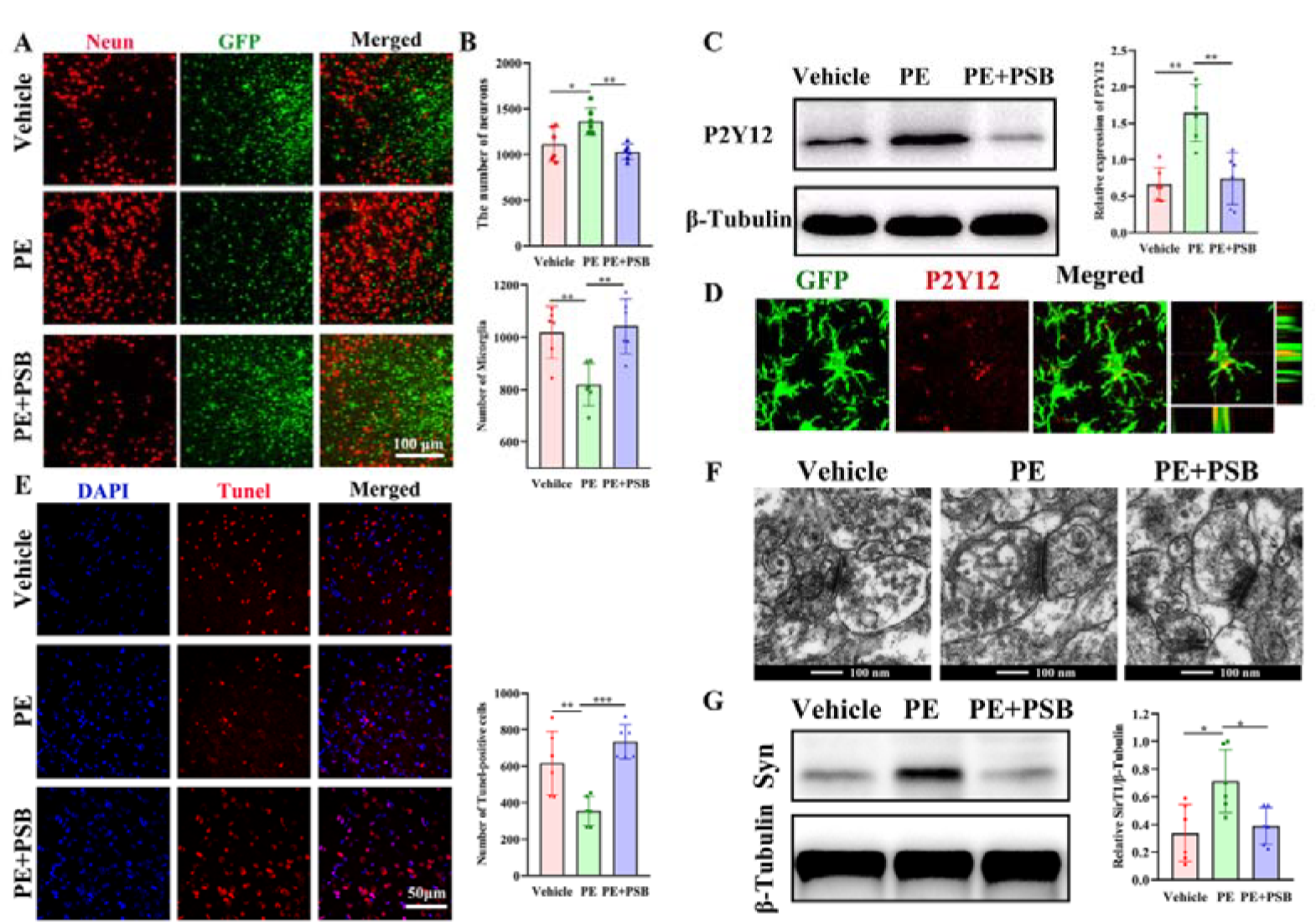
Physical exercise (PE) protected against brain damage after ischemic stroke and increased the expression of microglial P2Y12. A. Immunofluorescence staining of Neurons and GFP-positive microglia, ; B. Comparisons of neuronal and microglial numbers among the vehicle, PE, and PE+PSB0739 groups. C. Chemiluminescencec Comparison of the P2Y12 level among the vehicle, PE, and PE+PSB0739 groups. D. Immunofluorescence staining of P2Y12 in GFP-positive microglia. E. Comparisons of TUNEL-positive cells in peri-infarct areas among the vehicle, PE, and PE+PSB0739 groups. F. Representative images of the synaptic structures captured by transmission electron microscope (TEM) in vehicle, PE, and PE+PSB0739 groups. G. Comparisons of the synaptophysin (Syn) expression among the vehicle, PE, and PE+PSB0739 groups. Each dataset is expressed as the mean ± SD. \**P* ≤ 0.05; \*\**P* ≤ 0.01; \*\*\**P* ≤ 0.001; \*\*\*\**P* ≤ 0.0001. n = 6 mice.

### Endocannabinoid signaling was required for PE-regulated microglial dynamics nad microglia-neuron contacts

Endocannabinoid signaling (ECS) is comprised of 2-arachidonoylglycerol and anandamide (AEA), and their type-1 and type-2 cannabinoid receptors, CB1R and CB2R^[33]^, have been reported to be involved in neuron-microglia communication ^[18–21]^. To further explore the involvement of endocannabinoid system in PE-associated protection as well as in regulation of P2Y12 signaling, we detected the endocannabinoid receptors (CBRs) in brain. Western blotting showed that PE increased the expressions of both CBR1 (*P* < 0.01) and CBR2 (*P* < 0.01), when compared with that of the vehicle group (Fig. 4A). Further results of co-immunofluorescence staining showed that the up-regulated CBR1 were mainly located in neurons, not in the microglia (Fig. 4B); and the up-regulated CBR2 were mainly located in the microglia, not in neurons. We therefore treated PE animals with the CBR2 antagonist, AM630, to further characterize the mechanism of endocannabinoid (eCB) signaling in microglial extension and retraction, as well as during the microglia-neuron communication. We also treated stroke animals with URB597, the selective FAAH inhibitors to increase endocannabinoid signaling in brain^[34]^, to study whether the PE-related protection could be imitated (Fig. 4C). As expected, URB597 treatment significantly increased the expressions of CBR1 and CB2R, when compared with that of the vehicle group (*P* < 0.05 and *P* < 0.05, respectively), which did not differ from the PE group (both, *P* > 0.05) (Fig. 4D). AM630 treatment significantly decreased the expression of CBR2, when compared with that of the PE group (*P* < 0.05), while it did not affect the expression of CBR 1(*P* > 0.05; PE+AM630 *vs.* the PE group) (Fig. 4D). The expression of CB1R in the PE+AM630 group was higher than that in the vehicle group (*P* <0.05), but the expression of CBR2 did not differ from that in the vehicle group (*P* > 0.05). Additionally, URB597 treatment significantly increased the expression of P2Y12, compared with that of the vehicle group (*P* < 0.05), which did not differ from that of the PE group (*P* > 0.05) (Fig. 4E). The expression of P2Y12 was much less in the PE+AM630 group than that in the PE group (*P* < 0.05), which did not differ from that in the vehicle group (*P* > 0.05) (Fig. 4E). Immunofluorescence staining showed that URB597 treatment significantly increased the number of neurons (*P* < 0.01), while it decreased the number of microglia (*P* < 0.01) (Fig. 4F, G). The numbers of neurons and microglia in the URB597 group did not differ from those in the PE group (all, *P* > 0.05). However, when compared with the PE group, AM630 treatment significantly decreased the number of neurons in the peri-infarct areas (*P* < 0.05), while it increased the number of microglia (*P* < 0.05) (Fig. 4F, G).

**Figure 4.**
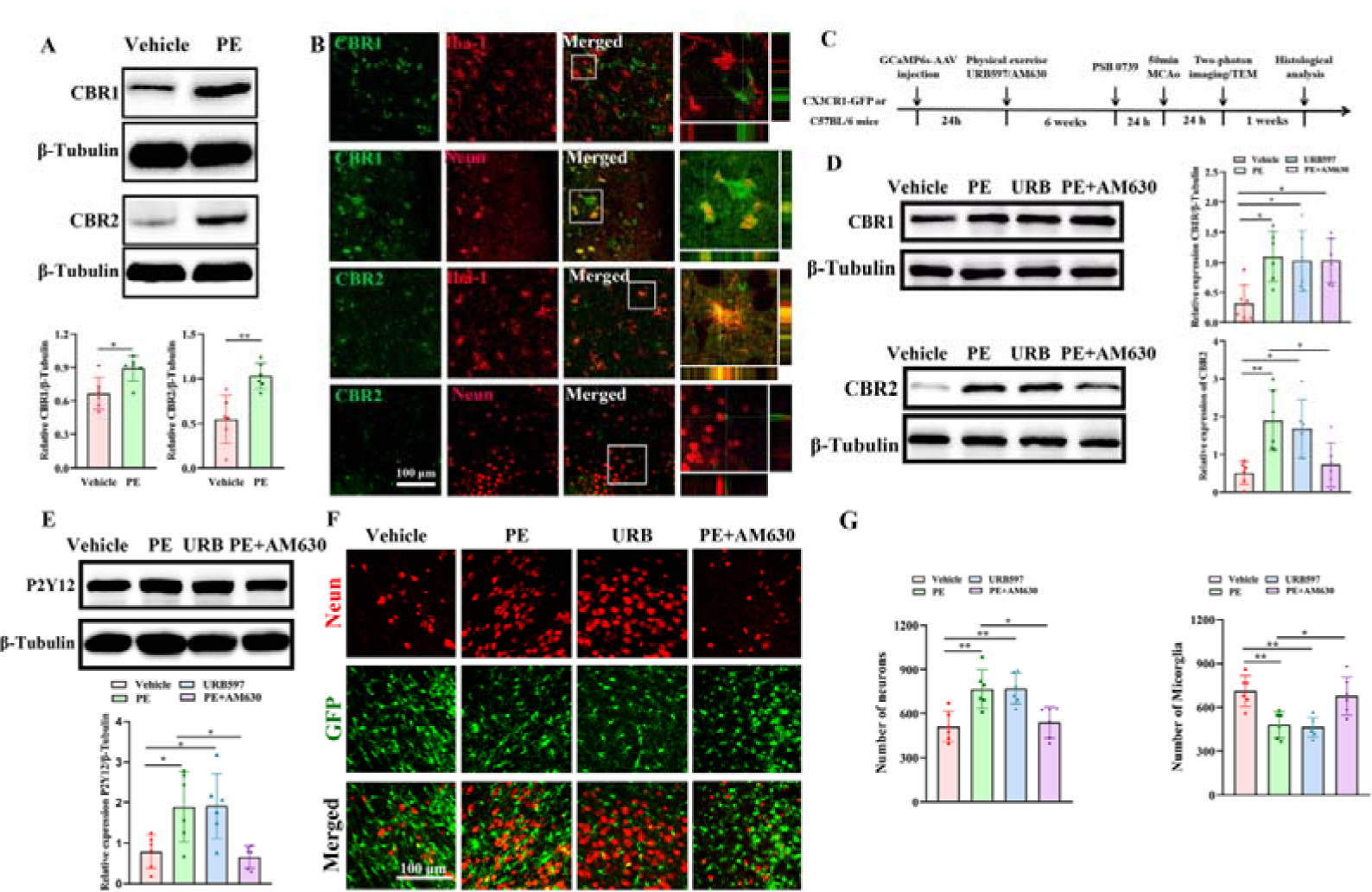
URB597 treatment mimicked the protective events induced by physical exercise (PE), while AM630 abolished the PE-associated protection. A. Comparisons of CBR1 and CBR2 expressions between the vehicle and PE groups. B. Immunofluorescent results of Neun, Iba-1, CBR1 and CBR2 showed that CBR1 was mainly located in neurons, CBR2 was mainly located in microglia. C. Schematic diagram showed the time points of URB597 or AM630 administration. D. Comparisons of the CBR1 and CBR2 expressions among the vehicle, PE, URB597 and PE+AM630 groups. E. Comparisons of P2Y12 expressions among the vehicle, PE, URB597 and PE+AM630 groups. F. Immunofluorescence staining of Neun using CX3CR1-GFP mice. G. Comparisons of Neurons and GFP-positive microglia among the vehicle, PE, URB597 and PE+AM630 groups. Each dataset is expressed as the mean ± SD. \**P* ≤ 0.05; \*\**P* ≤ 0.01; \*\*\**P* ≤ 0.001; \*\*\*\**P* ≤ 0.0001. n = 6 mice.

We next detected the involvement of endocannabinoid signaling in microglial processes as well as in PE-associated protection of neuronal activity using CXCR1-GFP mice with two-photon imaging, microglial motility was quantitated in the same cells for each group during a 2 min time period. Microglial branches were more pronounced in URB597 group (Supplementary vedio.4) than that in the vehicle group, which was similar to the PE groups. However, the vigorously microglial process disappeared in the PE+AM630 group (Fig. 5A) (Supplementary vedio. 5), the microglial images captured at 120s of scanning were overlaid with the same field that was captured at the start of scanning (Fig. 5B), which verified the dynamic state of microglia in the PE and URB597 groups. Specifically, two-way ANOVA results shows that the change ratios of the branch lengths in PE and URB597 groups were significantly increased than that in vehicle group at different time points, which were abolished in PE+AM630 group (Fig. 5C). Neuronal calcium activity was assessed by measuring the fluorescence intensity change of GCaMP6s, a high signal-to-noise and ultrasensitive fluorescent indicators detecting the increases in intracellular Ca^2+^ in a 1-min time period. Fig 6A shows that in the PE and URB597 groups, much microglial branches contacting with the dtomato-positive neurons in the vehicle and PE+AM630 groups, whereas fewer microglial branches contacting with the neurons. As expected, the amplitudes and frequencies of neuronal calcium in the URB597 group were significantly lower than those in the vehicle group (*P* < 0.001 and *P* < 0.01, respectively), which did not differ from the PE group (both *P* > 0.05). However, AM630 treatment significantly increased the amplitudes and frequencies of neuronal calcium activities, when compared with the PE group (*P* < 0.05 and *P* < 0.01, respectively), which did not differ from that of the vehicle group (all, *P* > 0.05) (Fig. 6A-C), indicating the microglial branches in the PE and URB597 groups may continuously monitor the neuronal calcium activities by contacting with the neuronal bodies. The TEM results verified the membrane structure of microglia-neuron contacts in PE and URB597 groups, but the membrane structure of microglia-neuron contacts did not observed in vehicle or PE+AM630 groups (Fig. 6D). PE and URB597 treatment kept the mitochondrial tubular networks and integrity, whereas which was swelled and cristae cracked. As expected, the western blotting (Fig. 6E) results verified that the expression of neuronal TOMM20 was significantly increased in the URB597 group, when compared with that of the vehicle group (*P* < 0.001), which did not differ from that of the PE group (*P* > 0.05) (Fig. 6E). However, AM630 treatment significantly decreased the expression of neuronal TOMM20 in PE mice (*P* < 0.05; PE+AM630 *vs.* PE groups), which did not differ from that of the vehicle group (*P* > 0.05, PE+AM630 *vs.* the vehicle groups) (Fig. 6F).

**Figure 6.**
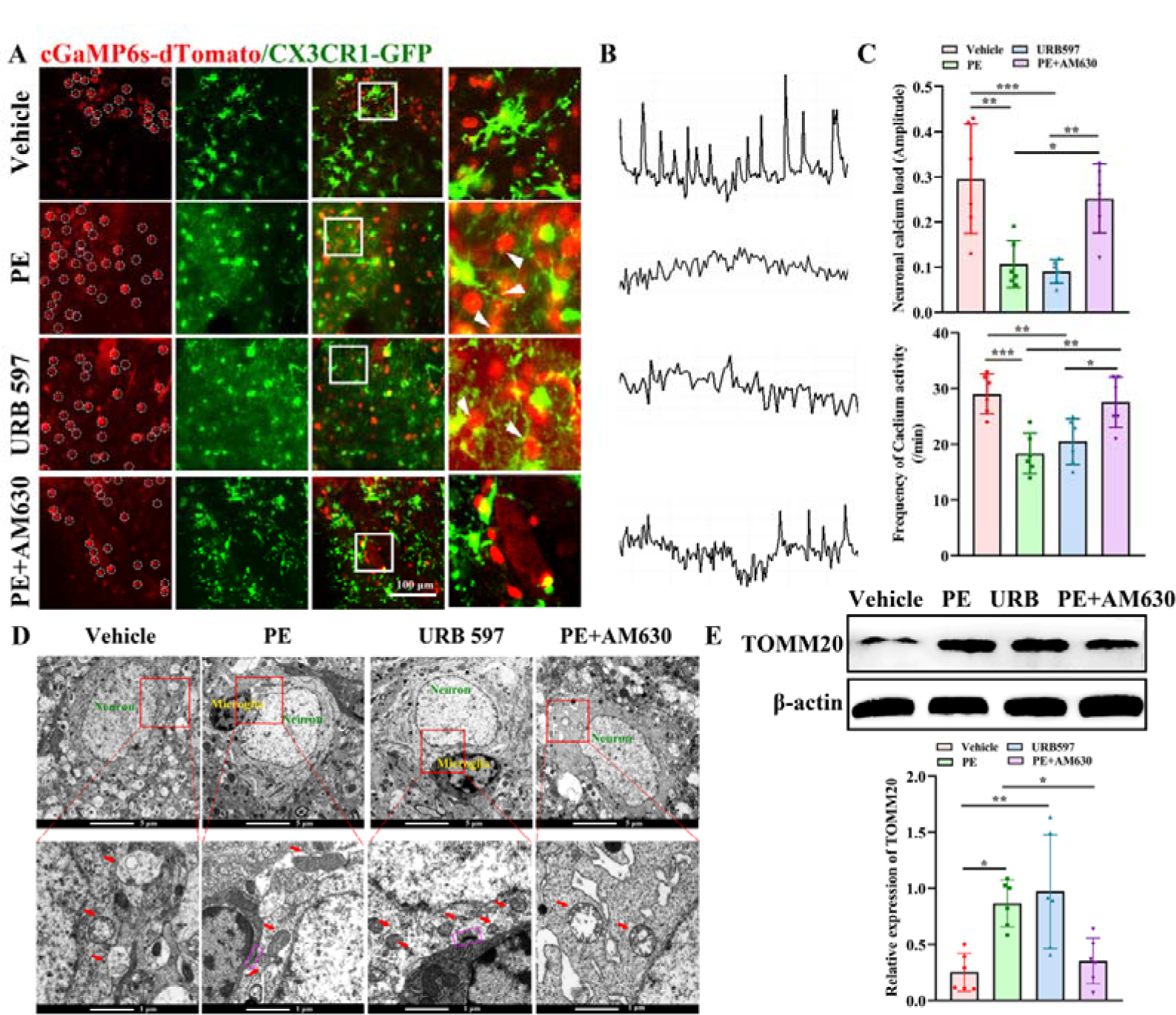
URB597 mimicked the improvement of microglial-neuron interactions induced by physical exercise, while AM630 decreased the protection of PE. A. Representative images showing GCaMP6s signal changes in neurons imaged at 24 h after transient middle cerebral artery occlusion.. B. Representative images about the immunofluorescent intensity changes of GCaMP6s-dTomato. C. Comparisons of the amplitudes and frequencies of the GCaMP6s signal changes among the vehicle, PE, URB597 and PE+AM630 groups. D. Representative images of the neuronal mitochondria and the microglia-neuron membrane junction captured by transmission electron microscope (TEM). Upper panel: images captured by1200X objective, red box labeled the magnified site showed in the below panel. below panel: images captured using 11000X objective, red arrows indicates the neuronal mitochondria and purple box in PE and URB597 group labeled the microglia-neuron junction. E. Comparison of TOM20 expression among the vehicle, PE, URB597 and PE+AM630 groups (Each dataset is expressed as the mean ± SD. \**P* ≤ 0.05; \*\**P* ≤ 0.01; \*\*\**P* ≤ 0.001; \*\*\*\**P* ≤ 0.0001. n = 6 mice)

URB597 significantly decreased the number of TUNEL-positive cells in peri-infarct areas (Fig. 7A-B), when compared with that in the vehicle group (*P* < 0.05), which did not differ from that of the PE group (*P* > 0.05). However, AM630 treatment significantly increased the number of TUNEL-positive cells, when compared with the PE group (*P* < 0.01), which did not differ from that of the vehicle group (*P* > 0.05). The TEM results show that the synaptic vesicles were much more in the PE and URP597 group compared to that in vehicle group, which was less in PE+AM630 group.The membrane structure of synapses were also clearly visible in PE and URB597 groups, whereas which was obscure in vehicle and PE+AM630 groups. Western blotting of synaptophysin (Syn) confirmed that URB597 significantly increased the relative expressions of Syn, when compared with the vehicle group (*P* < 0.0001), which did not differ from that of the PE group (*P* > 0.05) (Fig. 7D). However, injection of AM630 significantly decreased the relative expressions of Syn, when compared with that of the PE group (*P* < 0.0001), which did not differ from that of the vehicle group (*P* > 0.05) (Fig. 7D). Furthermore, URB597 treatment decreased the levels of pro-inflammatory cytokines including TNFα and iNOS, when compared with that in the vehicle group (*P* < 0.01, *P* < 0.001, respectively) (Fig. 7E, F), which did not differ from that in the PE group (both *P* > 0.05). However, the expressions of TNFα and iNOS in the PE+AM630 group were significantly increased, when compared with that in the PE group (*P* < 0.01, *P* < 0.0001, respectively), which did not differ from that in the vehicle group (both *P* > 0.05). In contrast, URB597 treatment increased IL-10 levels, when compared with that in the vehicle group (*P* < 0.01), which did not differ from that in the PE group (*P* > 0.05) (Fig. 7G),. However, the level of IL-10 was significantly decreased in the PE+AM630 group, when compared with that in the PE group (*P* < 0.0001), which did not differ from that in the vehicle group (*P* > 0.05) (Fig. 7G),. Together, these results showed that URB597 imitated the effect of microglial dynamics induced by PE, and enhanced microglial motility and inhibited microglial neuroinflammation. Furthermore, AM630 blocked PE-associated events, indicating that PE promoted microglial processes and inhibited neuroinflammation via endocannabinoid signaling.

**Figure 7.**
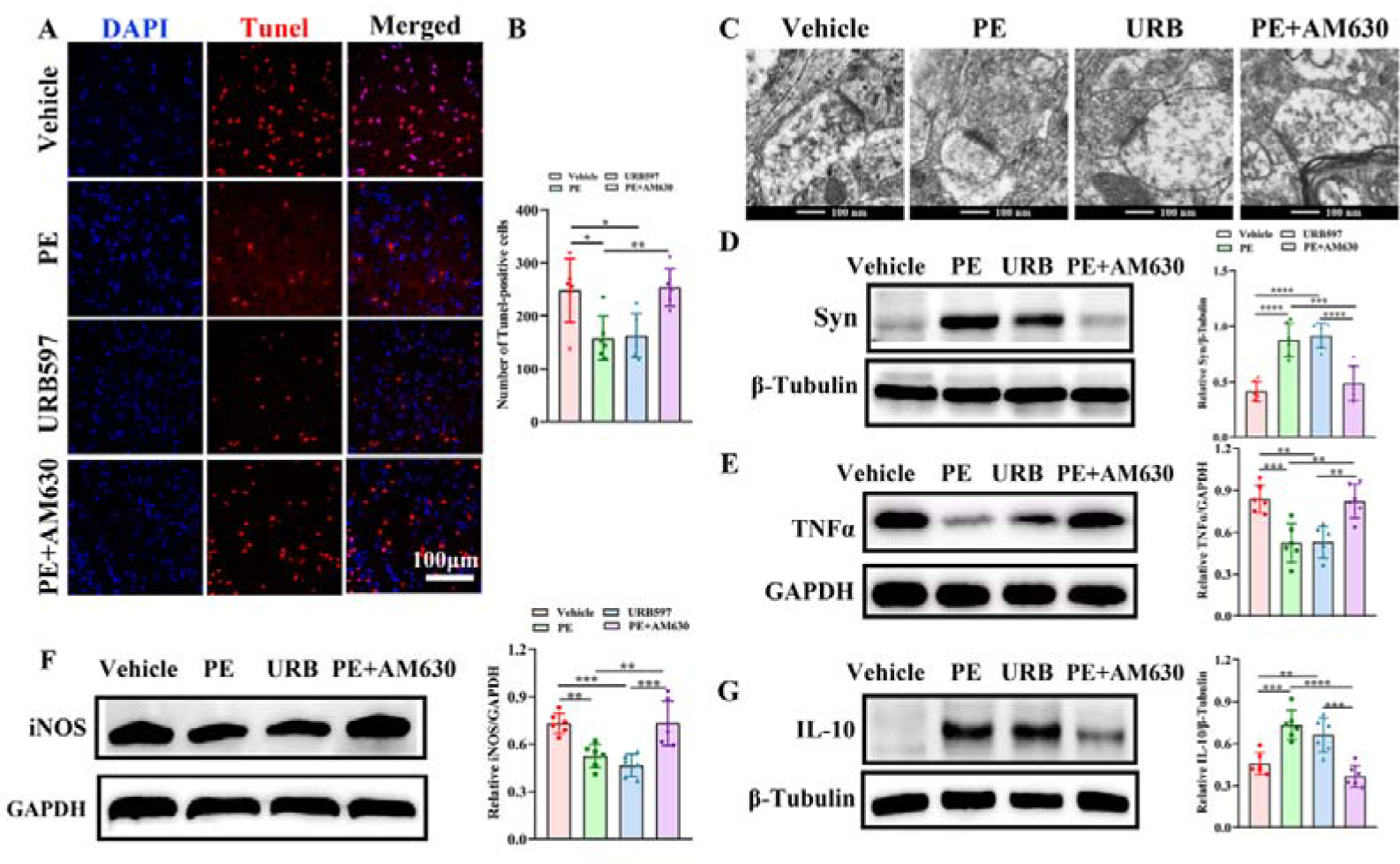
URB597 and PE decreased neuronal apoptosis, protected against the synaptic damage and decreased the pro-inflammatory response, whereas AM630 restrained the physical exercise (PE)-associated effects. A. Representative images for TUNEL-positive cells in the peri-infarct areas. B. Comparison of TUNEL-positive cell numbers among the vehicle, PE, URB597 and PE+AM630 groups. C. Representative images of the synaptic structures captured by transmission electron microscope (TEM) in vehicle, PE, and PE+PSB0739 groups. D. Comparison of synaptophysin (Syn) expression among the vehicle, PE, URB597 and PE+AM630 groups. E. Comparison of TNFα expression among the vehicle, PE, URB597 and PE+AM630 groups. F. Comparison of iNOS expression among the vehicle, PE, URB597 and PE+AM630 groups. G. Comparison of IL-10 expression among the vehicle, PE, URB597 and PE+AM630 groups (Each dataset is expressed as the mean ± SD. \**P* ≤ 0.05; \*\**P* ≤ 0.01; \*\*\**P* ≤ 0.001; \*\*\*\**P* ≤ 0.0001. n = 6 mice).

### Endocannabinoid system may act as the upstream of P2Y12 signaling

To further characterize the interaction of endocannabinoid signaling and microglial P2Y12 in PE-associated protection, we injected PSB0739 followed by the URB597 treatment before the induction of stroke surgery (URB597+PSB0739 group) (Fig.8A). URB597 increased the expression of P2Y12 (*P* < 0.05), which was remarkably abolished by PSB0739 (*P* < 0.05), which showed no significant difference in URB597+PSB0739 group compared to that in vehicle group (*P* > 0.5) (Fig.8B). However, PSB0739 injection did not affect the expressions of CBR1 and CBR2 upregulated by URB597 (*P* > 0.05) (Fig. 8C-D). Additionally, URB597 significantly decreased the expressions of pro-inflammatory cytokines including iNOS and TNFα, when compared with that of the vehicle group (both *P* < 0.01), while it did not differ in the URB597+PSB0739 group (both *P* > 0.05). Expressions of these pro-inflammatory cytokines were much higher in the URB597+PSB0739 group than that in the URB597 group (*P* < 0.01, *P* < 0.05, respectively) (Fig. 8E). In contrast, URB597 treatment increased the level of IL-10, when compared with that of the vehicle group (*P* < 0.05), and showed no difference when compared with the URB597+PSB0739 group (*P* > 0.05). The IL-10 level was significantly decreased in the URB597+PSB0739 group, when compared with that of the URB597 group (*P* < 0.05) (Fig. 8F).

**Figure 8.**
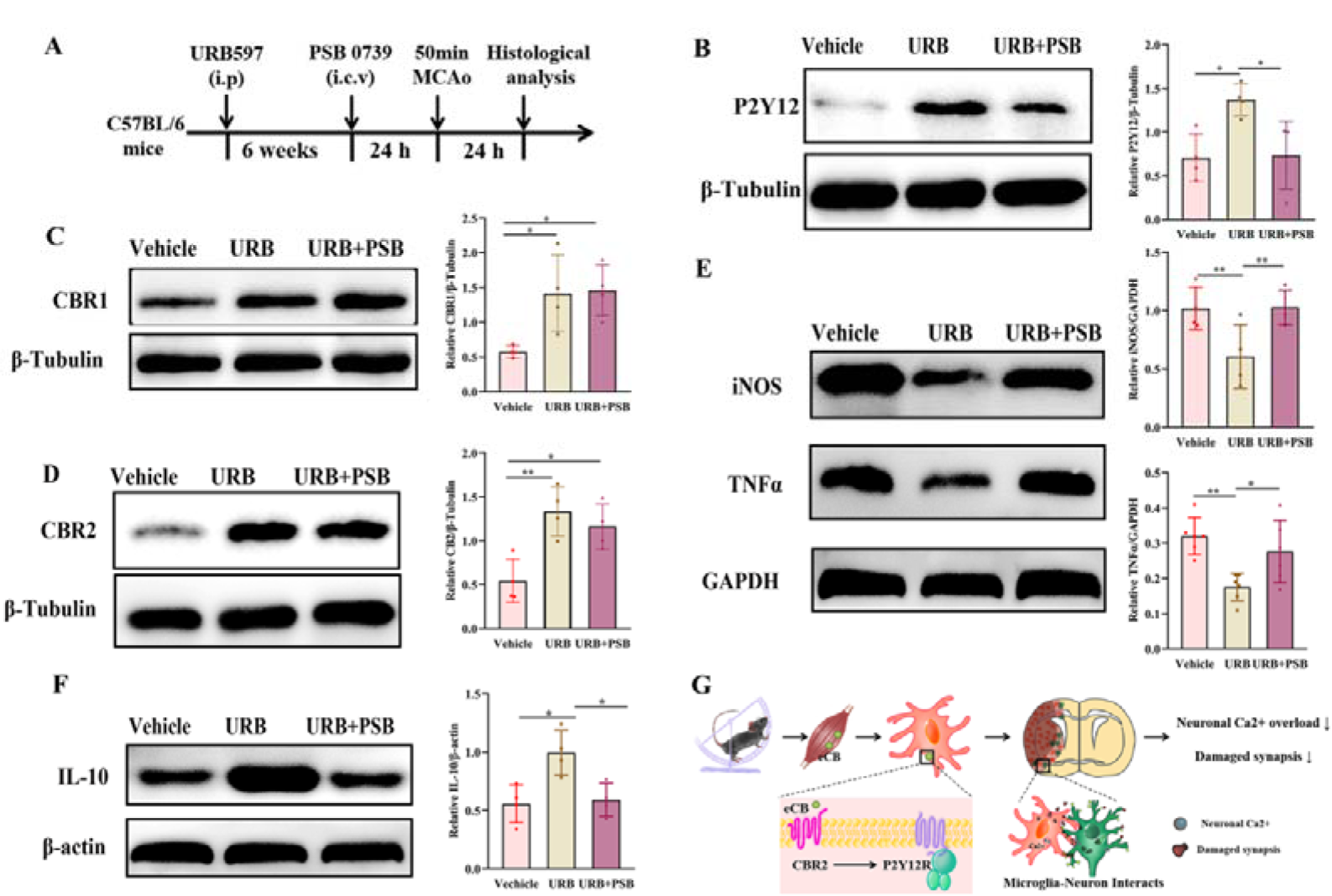
The endocannabinoid system acts as the upstream of P2Y12 signaling. A. Schematic diagram showed the time points of URB597 and PSB0739 administration. B. Comparison of P2Y12 expression among the vehicle, URB597 and URB597+ PSB0739 groups. C. Comparison of CBR1 expression among the vehicle, URB597 and URB597+ PSB0739 groups. D. Comparison of CBR2 expression among the vehicle, URB597 and URB597+ PSB0739 groups. E. Comparison of iNOS, TNF α expressions among the vehicle, URB597 and URB597+ PSB0739 groups. F. Comparison of IL-10 expression among the vehicle, URB597 and URB597+ PSB0739 groups. G. The hypothesis diagram of this study (Each dataset is expressed as the mean ± SD. \**P* ≤ 0.05; \*\**P* ≤ 0.01; \*\*\**P* ≤ 0.001; \*\*\*\**P* ≤ 0.0001. n = 4 mice).

Collectively, these results suggested that URB597 treatment imitated PE-related protection by upregulating the expressions of CBR1 and CBR2, increasing P2Y12 levels, and inhibiting neuroinflammation, to further promote neuronal survival in peri-infarct areas. However, AM630 treatment abolished PE-associated protection, it neutralized the upregulated expressions of CBR2 and P2Y12, increased microglial activation, and promoted neuronal death. Additionally, our result also showed that blockade of P2Y12 did not affect the expressions of CBR1 and CBR2 upregulated by URB597, it indicates that CBR2 is the upstream of P2Y12 signaling regulating the microglial dynamics and the microglia-neuron communication (Fig. 8F).

## Discussion

There is strong evidence that microglia represent the first line of defense against ischemic stroke, which leads to either pro- or anti-inflammatory phenotypes^[35,36]^. However, recent reports supported the idea that two phenotypes of microglia co-exist in single microglia, rather than an “all or none” phenomenon. They also overlap during disease progression and may therefore antagonize each other^[35]^. Previous studies have reported that depletion of microglia led to calcium overload, increasing post-ischemic inflammation and worsening ischemic brain damage^[28,35,37]^, further suggesting that microglia were needed to reduce ischemic injury. Importantly, neuronal hyperactivities or hypoactivities induced microglial extension and outgrowth, further interacting with neuronal somata and dendrites^[10, 38, 39]^, and the durations of microglial-neuronal interactions were prolonged after transient cerebral ischemia^[40]^, suggesting the protective role of microglia. Consistently, using *in vivo* two-photon time-lapse imaging, we found that microglia grew and retracted their branches when dynamically contacting neurons, which contributed to calcium homeostasis and protected against the dying of neurons in penumbra areas during the acute stages of ischemic stroke. We specifically showed that PE promoted microglial processes and increased microglial-neuronal junctions, to further inhibit calcium overloading, and increase the levels of anti-inflammatory cytokines to eventually improve neuronal functioning.

We have demonstrated that PE protected against neuronal detachment, by reducing the calcium overload, which was correlated with excitotoxicity leading to neuronal death^[41]^. Microglia-neuronal junctions are responsible for calcium homeostasis during brain damage^[10]^, and elimination of microglia or blockade of microglia-neuronal junctions both lead to neuronal calcium overload and increased neuronal death^[10, 41]^. PE enhanced the microglial outgrowth process and increased microglial-neuronal junctions during stroke. Furthermore, PE decreased levels of pro-inflammatory cytokines by inhibiting neuronal calcium overloading, because the immune response to ischemic injury has been reported to peak later than calcium overloading^[35,42]^. An interesting question is how microglial processes inhibit the neuronal calcium overload. One reasonable mechanism may be attributable to the phagocytic activity of microglia in the presence of cytosolic calcium. The majority of microglia are at a midpoint between surveying and being fully activated, so they can effectively phagocytize their targets without becoming activated^[43]^. Consistent with this possibility, microglia-neuron junctions have been shown to have an increased number of lysosomes^[10]^, and hyperactive neurons have been shown to increase the calcium influx of microglia^[38]^.

The mechanisms involving PE regulating microglial processes and microglial-neuron interactions have not been clearly identified. Previous evidence has shown that P2Y12 canonically drives both the microglial outgrowth process^[44]^ as well as formation of microglia-neuron junctions^[10]^, contributing to microglial protection. P2Y12R is the primary site at which nucleotides act to induce microglial motility and mediate the microglial chemotaxis process towards the “find-me” signal, ADP/ATP gradients reported during the early stages of brain injury^[45,46]^, which are dramatically reduced after microglial activation^[47]^. Furthermore, microglial process numbers and somatic microglia-neuron junctions^[10,44]^ decreased in the absence of P2Y12^[10]^. Consistent with this possibility, we showed that PE inhibited microglial activation and increased the levels of microglial P2Y12, to further enhance microglial dynamics and increase the number of microglial-neuronal junctions, while blockade of P2Y12 neutralized the exercise-mediated microglial response and increased post-stroke injury.

ECS represents a newly discovered mechanism of neuro-immune communication during brain development or brain injury^[48,49]^. We have identified ECS as the link between physical exercise and microglial dynamics, as well as for microglia-neuron communication. ECS has been shown to regulate transcriptional programs in microglia^[50]^, and exercise increases microglial IL-10 expression in an ECS-dependent manner^[51]^. ECS involves the CBR1 and CBR2, which are expressed in neurons and microglia, respectively^[21, 43]^. We verified that PE increased the expression of microglial P2Y12 through the CBR2. Specifically, exercise and URB597 treatment both increased the levels of CBR2, upregulated P2Y12 expression, and improved microglial dynamics, whereas the CBR2 antagonist abolished the increase of P2Y12 induced by PE, counteracting its protective events. It is worth noting that blocking of CBR2 abolished exercise-induced upregulation of P2Y12, while PSB0739 did not affect the expression of CBRs, suggesting the predominance of ECS.

P2Y12 upregulated by ECS is of special importance because it is also sensitive to changes in the levels of γ-aminobutyric acid (GABA)^[52]^, which mediates microglial chemotaxis and motility in damaged areas^[53]^. Notably, ECS bidirectionally regulates the interactions between neurons and microglia. ECS modulates the secretion of neurotransmitters from neurons that may affect microglia^[54]^, so it is possible that PE increases the expression of CBR1 in neurons via CBR2 in microglia, because the majority of CBR1 are located on GABAergic neurons. ECS provides a feedback mechanism by which microglia alter the endocannabinoid response to neuronal CBR1, to mediate GABAergic signaling^[55]^. Microglial processes are also required for remodeling of neuronal circuits after brain injury, because they monitor synaptic functioning by interacting with axon buttons or dendritic spines ^[56]^, which may restore synapse function in the ischemic brain^[40]^. Indeed, our results showed that exercise increased synaptic vesicles and synaptophysin expressions.

## Data availability

The datasets analyzed in this study are available from the corresponding author upon reasonable request. Source data are provided with this paper.

## Abbreviations

physical exercise:PE; transit middle cerebral artery occlusion:tMCAo; adenosine diphosphate: ADP; G protein-coupled: GPCR; Anandamide: AEA; Cannabinoid 1 receptor:CB1R; Cannabinoid 2 receptor:CB2R; fatty acid amide hydrolase:FAAH; calcium:Ca2+; endocannabinoid signaling: ECS; adeno-associated virus: AAV; Ionized calcium-binding adapter molecule 1: Iba 1.

## Acknowledgements

We thank Neurology Department in the First Affifiliated Hospital of Sun Yat-sen University (Guangdong Provincial Engineering Center for Major Neurological Disease Treatment; Guangdong Provincial Translational Medicine Innovation Platform for Diagnosis and Treatment of Major Neurological Disease) for providing the experimental equipment, and Guangdong Laboratory Animals Monitoring Institute for breeding the CX3CR1-GFP mice.

## Authors’ contributions

Xiao-fei He performed the Two-photon imaging; Ge Li bred the CX3CR1-GFP mice, Xiao-fei He, Yun Zhao, Jing-hui Xu and Jing Luo performed the Histological staining and Western blotting; Xiao-fei He and Yun Zhao draft the manuscript; Hai-qing Zheng, Li-ying Zhang and Xi-quan Hu revised the manuscript. Li-ying Zhang and Xi-quan Hu conceived and designed the research, and edited and revised the manuscript. The authors read and approved the final manuscript.

